# Locally heterogeneous soil viral and prokaryotic responses to prescribed burn correspond with patchy burn severity in a mixed conifer forest

**DOI:** 10.1101/2024.10.28.619755

**Authors:** Sara E. Geonczy, Luke S. Hillary, Christian Santos-Medellín, Jess W. Sorensen, Joanne B. Emerson

## Abstract

Prescribed burning, a strategy to mitigate wildfires, imparts physicochemical and biological changes to soil. The effects of burns on soil viruses and virus-host dynamics are largely unexplored, despite known viral and prokaryotic contributions to biogeochemical processes. Using a viromic (<0.2 µm size fraction metagenomic) approach, we assessed how viral communities responded to a spring prescribed burn in a mixed conifer forest and whether soil chemical properties and/or prokaryotic host communities could explain the observed patterns. From 120 soil samples (two per depth at 0-3 and 3-6 cm from four burned and two control plots at five timepoints, two before and three after the burn), 91 viromes and 115 16S rRNA gene amplicon libraries were sequenced. Plot location had the greatest effect on explaining variance in viral communities, over treatment (burned or not), depth, and timepoint. Viral and prokaryotic communities exhibited locally heterogenous responses to the fire, with some burned communities resembling unburned controls. This was attributed to patchy burn severity (defined by soil chemistry). Low viromic DNA yields indicated substantial loss of viral biomass in high-severity locations. The relative abundances of Firmicutes, Actinobacteria, and the viruses predicted to infect them significantly increased along the burn severity gradient, suggesting survival of spore formers and viral infection of these abundant, fire-responsive taxa. Our analyses highlight the importance of a nuanced view of soil community responses to fire, not just to burn overall, but to the specific degree of burn severity experienced by each patch of soil, which differed for nearby soils in the same fire.

## INTRODUCTION

Prescribed burning is a management practice that aims to reintroduce fire as an ecological process in areas where fire has otherwise been suppressed, while at the same time aiding in the prevention of catastrophic wildfires (1–5). This practice is one aspect of regulating forest biogeochemical cycling (6), given that forests are a substantial carbon sink (7), but disturbances like wildfires can make forests a potential carbon source (8). Naturally, there is interest in the impact of prescribed burning on soil microorganisms, which are integral to key soil processes that mediate carbon turnover and nutrient availability in forest ecosystems (9–11). While most studies have focused on fungi and bacteria, for example identifying pyrophilous taxa that positively respond to fire (12–14), the impact of prescribed burning on abundant soil viruses has only recently been considered and is largely unexplored (15).

Viruses apply selective pressure on cellular organisms via infection and lysis and may also have distinct impacts on biogeochemical cycling (16). For example, in marine ecosystems, viruses are known to affect biogeochemical cycling through conversion of living microbial biomass into dissolved organic matter, contributing to an estimated 10-40% of total bacterial mortality daily, including 10% of phytoplankton (17). Incorporating soil viruses into the microbial community framework has begun to elucidate their role in key soil processes (18), which would presumably be impacted by disturbances like fire.

In response to varying degrees of fire, including prescribed burning, there can be significant effects on the biomass, abundance, richness, evenness, and diversity of soil biota across many ecosystems (12,13,19,20). Prescribed burning in forests typically reaches higher temperatures than in other ecosystems (21), and heating directly impacts the survival of soil biota (22). Changes to soil properties, including short-term reductions in organic matter and increases in pH, mineral phosphorus, total nitrogen, potassium, calcium, and magnesium can also impact microbial communities (22–24). These changes are especially evident at the soil surface (25,26), as heat penetration is typically only surface-level, down to 2-3 cm, depending on soil moisture, texture, and compaction, and the nature of the fire itself (27,28). Post-fire community changes show distinct microbial zones along a soil profile (29), indicating the importance of depth-resolved studies in soils post-fire. Viruses are known to be sensitive to heating (30–32). However, fire has unique heating behavior and can impact soil properties due to fuel combustion (26), and these complexities are not captured when studying the effect of temperature change alone. Studies have begun to explore the effect of fire on soil viral communities (33), but questions remain in understanding whether viruses and their hosts exhibit similar survival patterns, and to what extent infection dynamics play a role in community compositional changes, in the short-term period after a fire.

Burn severity, the degree of fire-induced change to soil (34), is a factor of fire intensity and duration (22) and is an important consideration for understanding fire’s effects on microbial communities (13,35). However, the characteristic heterogeneity of burn severity across the landscape can be difficult to capture at a small scale relevant to microorganisms (20). Prescribed fires are designed to be of low severity and are considered effective at reducing surface fuel with low tree mortality (36). While summer and fall burns are more congruent with the natural timing of fires as an ecosystem process, changing environmental conditions have shortened the window for safe and effective burns even in the fall, with increasing consideration of winter and spring prescribed burns (37,38). These early season burns, when fuel moisture and soil moisture levels are higher, are less effective at consuming fuel and tend to have a greater degree of patchiness than late season burns (39), and therefore they may confer greater ecosystem resilience due to unburned refugia (5). Understanding microbial community responses to these prescribed burns therefore has practical applications, both for future prescribed burns under similar conditions and because these burns offer more controlled, safe, and tractable study designs (e.g., including pre-fire sampling opportunities) than wildfires, while still in a field-scale setting.

Here, we investigated the impact of prescribed burning on dsDNA viral communities in a forest soil in response to a spring prescribed burn, while simultaneously considering the burn impact on putative prokaryotic hosts (via 16S rRNA gene amplicon sequencing) and soil chemical properties. Leveraging a viromic (< 0.2 µm viral size fraction metagenomic) approach (40), we assessed how viral communities changed in response to the prescribed burn, whether the response differed between two near-surface depths (0-3 cm and 3-6 cm), and to what extent soil chemical properties and/or microbial community composition (including specific taxa predicted to be preyed upon by the most abundant viruses) could explain the observed patterns.

## MATERIALS AND METHODS

### Study site description

We set up six 5 x 5 m square plots across two 20-acre forest units at the Blodgett Forest Research Station in California, USA (Figure 1, GPS coordinates in **Supplementary Table 1**). The two units had similar vegetation and management history, including over 100 years of fire exclusion (41). Three plots were in a forest unit that was planned to burn in a spring prescribed burn. The other three plots were in a unit that was not planned to be burned, and thus these plots were initially selected to represent control plots, but one plot in the control area burned (see “Prescribed Burn Treatment” section for further details). There was an attempt to locate plots approximately the same distance away from each other in each unit but based on our plot criteria (similar characteristics in terms of litter quality and quantity, slope, and without trees or shrubs directly growing in the plot), our choices were constrained, and the distance between plots varied. Within each forest unit, the distance between plots was between 63 - 148 m in the burn unit and 89 - 271 m in the control unit. Between both units, the maximum distance between plots was 558 m, and the minimum distance was 283 m. All plots were located in the Holland-Musick loams, which consisted of fine-loamy soil with extensive profile development and an O horizon that extended from the surface to 5 to 8 cm deep (42). The vegetation surrounding all plots is considered mixed-conifer and includes a mix of firs (predominantly Douglas fir), pines (predominantly ponderosa), incense cedar, and oaks (tan oak and black oak).

**Figure 1.**
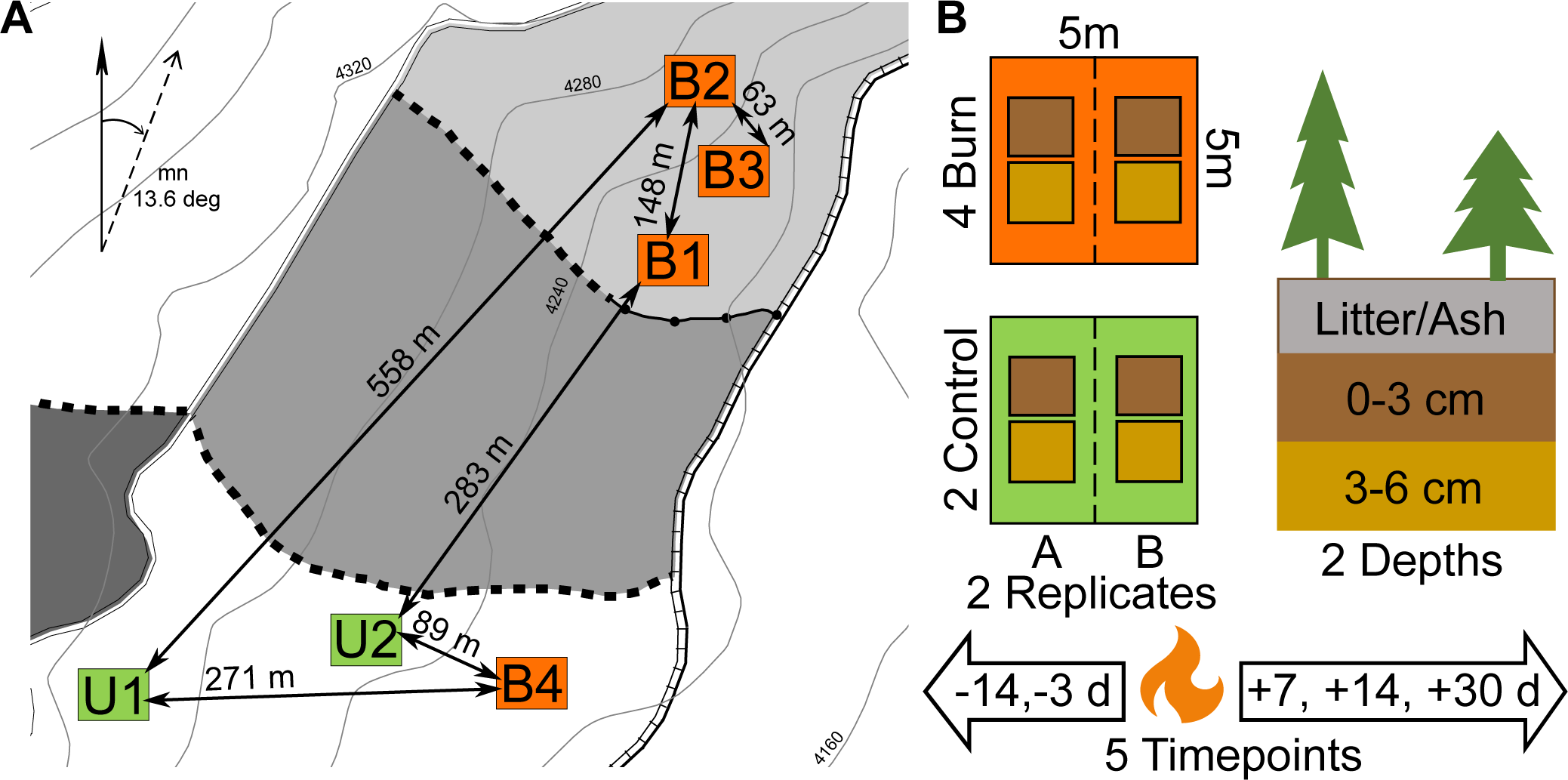
Experimental design overview, **A.** Map of plot locations with minimum and maximum distances indicated within units (plot groups) and between units. Orange squares indicate burn treatment plots and green squares indicate control plots. The vertical arrow represents true north and the angled arrow represents magnetic north, with a magnetic declination of 13.6 degrees. **B.** Schematic overview, indicating plot size and sampling scheme (4 burn treatment plots and 2 control plots sampled at 2 depths with 2 replicates per plot for five timepoints: two before the fire and three after the fire). Depth colors on the left correspond with depths in the plot diagrams on the right.

### Prescribed burn treatment

Two days prior to the prescribed burn (2021 April 19), we buried two data logging units 30 cm below the surface (Extech SDL200 4-Channel Data Logging Thermometer, Waltham, MA, USA) just outside each burn plot on opposite sides and one data logging unit just outside one side of each control plot. The data logging units had temperature probes (Extech TP873 Bead Wire Type K Temperature Probe, -30 to 300°C) inserted at 1, 2, 4, and 6 cm depth in the soil profile. These data logging units tracked soil temperature over six days, capturing temperature fluctuations before, during, and after the prescribed burn (**Supplementary Table 2**). One of the dataloggers for plot B3 completely burned, and one datalogger for B2 had an SD card that was corrupted during the burn, so we were only able to retrieve one set of temperature data (from one datalogger each) for these two plots.

Although the unit where our control plots were located was not planned to be burned, to determine whether the conditions were appropriate for a prescribed burn in the treatment unit, a test burn was conducted on 12 April 2021 in the unit with the control plots. To protect the control plots, a fire line was cut around the perimeter of each plot to exclude fire, with a 5-10 foot buffer from the edge of the plots. Additionally, there was a person with water at each plot to monitor and ensure fire did not enter any control plot during the burn. Unfortunately, one of the control plots burned as a result of this test burn, due to spread of the test fire after monitoring had been completed, (fire entered the plot on 23 April 2021 as evident from the temperature data) leaving only two control plots instead of the planned three for post-burn control samples. This fire behavior was consistent with the official prescribed burn, where spot fires were found over one month after the burn was set, marking an unusual season for prescribed burning.

The prescribed burn of the unit where the treatment plots were located began on 21 April 2021. It was considered a first entry spring burn that primarily used a strip head ignition pattern (linear strips of fire ignited on the ground evenly and closely together with the intention of homogenizing fire behavior and effects (5)). No ignitions occurred within the treatment plots, and fire was allowed to move into the plots from outside the plot boundaries. The fire was maintained as a surface fire with limited individual or small group tree torching. Smoldering, creeping, and snag and stumphole burning continued overnight and led to reignitions in later days, all within burn boundaries. Ignitions of both units (the test burn and the official prescribed burn) occurred during the same burn window (time period with acceptable burning conditions) under similar weather conditions, resulting in similar fire behavior and effects across units. This reinforced our decision to consider samples collected from the burned ‘control’ plot in the same treatment group as the other three burn plots.

### Soil sampling

Each plot was divided into two halves (subplots) along the middle so that we could sample two replicates in each plot (one from each subplot) at each of five timepoints (14 and 3 days before the burn and 7, 14, and 30 days after the burn). Each subplot was then divided into eight equal-sized square sampling units (two columns of four sampling units in each subplot). At each timepoint over the course of the study, we took soil samples in the middle of corresponding sampling units of each plot (e.g. the top right of the right subplot and the top right of the left subplot for a single timepoint). The distance between replicate samples within each plot was approximately 3.75 m. At each sampling location, we collected approximately 300 g of soil at two depths, between 0 and 3 cm and between 3 and 6 cm, for a total of 4 samples per timepoint per plot. We homogenized and sieved (8 mm) all samples separately, yielding a total of 24 samples for each timepoint and a total of 120 samples for the entire study. The samples were stored on ice and transported back to the lab the same day as collection and promptly placed at 4°C until further sample processing, which started within 48 hours.

### Field metadata collection

We inventoried the trees and shrubs surrounding each plot and collected a representative sample of litter to determine the mix of vegetation contributing to litter quality and quantity (**Supplementary Table 1**). During sample collection, in-field soil temperature measurements were taken for each depth at each sampling location. We tested water repellency by wetting a sample with a single drop of deionized water and counting how many seconds it took until the drop had fully infiltrated into the soil using a stopwatch (43). We also measured litter depth (or ash depth if it was post-burn) and O horizon depth (**Supplementary Table 3**). Additionally, we have precipitation data for March to June 2021 (**Supplementary Table 4**).

### Soil chemical measurements

We calculated the gravimetric soil moisture of each sample by weighing out 10 g of soil into an aluminum weigh dish and measuring the weight after 24 hours in a drying oven at 105°C. A suite of soil chemistry analyses under the Haney Soil Health Test package (44,45) was performed by Ward Laboratories (Kearney, Nebraska, USA) (**Supplementary Table 5**). For the first sampling timepoint, an equal mass of the replicates of each depth in each plot were combined, and 200 g of each homogenized field-condition soil (not dried) were submitted to determine average measurements for both samples, while for all subsequent timepoints, 200 g of field-condition soil of each individual sample were submitted for separate measurements. The measurements included: soil pH and soluble salts (1:1 soil/water suspension), organic matter (percentage weight loss-on-ignition), soil respiration (Solvita® CO_2_-Burst method: soil was dried at 50°C and sieved at 2 mm, then 40 g soil was incubated for 24 hours in a 50 mL container with perforated bottom allowing for rewetting with 25 mL distilled water in a 250 mL glass jar sealed with a Solvita® paddle, and after 24 hours, CO_2_-C was quantified with a Solvita® digital reader with the units of mg CO_2_-C/kg soil), water extractable total N and organic C (quantified by shaking 4 g of soil with 40 mL of distilled water for 10 min and filtering with a Whatman 2V), Haney-(H3A: combination of citric, malic, and oxalic acids) extracted ammonium (NH_4_-N), nitrate (NO_3_-N), and inorganic phosphorus (orthophosphate: PO_4_-P) (the latter quantified with flow injection analysis), and H3A-extracted total phosphorus (P), potassium (K), zinc (Zn), iron (Fe), manganese (Mn), copper (Cu), sulfur (S), calcium (Ca), magnesium (Mg), sodium (Na), and Aluminum (Al) (quantified using inductively coupled argon plasma, or ICAP, atomic emission spectroscopy). Calculations included inorganic N (sum of H3A-extracted NH_4_-N and NO_3_-N), organic N (water-extractable total N minus inorganic N), organic P (total P minus inorganic P), microbially active carbon (soil respiration divided by water extractable organic C, expressed as percentage), organic C : organic N, and organic N : inorganic N.

### Viromic DNA extraction and quantification

Within 72 hours of sample collection, we extracted viromic DNA from all samples (of the 24 samples per timepoint, 12 were randomly selected for processing each day, starting the day after collection), as previously described (32,46). Briefly, to 10 g of each soil sample, we added 9 mL of protein-supplemented phosphate-buffered saline solution (PPBS: 2% bovine serum albumin, 10% phosphate-buffered saline, 1% potassium citrate, and 150 mM MgSO4, pH 6.5). Each sample was vortexed until homogenized and then placed on an orbital shaker at 300 rpm at 4°C for 10 min.

Supernatant was transferred into a new tube, and then 9 mL of PPBS was added to the soil remaining in the original tube, repeating the process for a total of three soil resuspensions and elutions, ending with a total of three volumes of supernatant pooled together in a new centrifuge tube. For each sample, the pooled supernatant was then centrifuged for 8 min at 10,000 *g* at 4°C, collecting the progressively clearer supernatant and discarding the concentrated soil particles, for a total of three centrifugations and supernatant collections. The final supernatant was first filtered through a 5 µm polyethersulfone (PES) membrane filter (PALL, Port Washington, NY, USA) to filter out larger soil particles, and then the filtrate went through a 0.22 µm PES membrane filter (PALL) to exclude most cellular organisms (e.g., some ultra-small bacteria are known to be retained in this size fraction (47,48)). Viral-sized particles and DNA remaining in the filtrate were concentrated into a pellet using an Optima LE-80K ultracentrifuge with a 50.2 Ti rotor (Beckman-Coulter, Brea, CA, USA) at 111,818 *g* at 4°C for 2 h 25 min. The pellet was resuspended in 200 µL of 0.02 µm filtered ultrapure water and treated with DNase, which included an incubation at 37°C for 30 min after adding 10 µL of RQ1 RNase-free DNase (Promega, Madison, WI, USA). We then extracted DNA using the DNeasy PowerSoil Pro kit (Qiagen, Hilden, Germany), following the manufacturer’s protocol, with the addition of a 10 min incubation at 65°C after adding the lysis buffer and prior to the bead-beating step, which also served as the ‘stop’ incubation for the DNase treatment. DNA quantification was performed using the Qubit dsDNA HS Assay and Qubit 4 fluorometer (ThermoFisher, Waltham, MA, USA). For timepoints 1-4, samples with yields above 0.05 ng/µl or 0.49 ng/g soil (above detection limits) were prepared for library construction and sequencing. For timepoint 5 (30 days after the burn), viromic DNA yields for 10 of the 24 samples (eight of the 16 burned samples and two of the 8 control samples) were below detection limits, so the decision was made not to sequence any of the viromes for that timepoint.

### Total DNA extraction and quantification

For each sample, 0.25 g of soil was added directly to the DNeasy PowerSoil Pro kit (Qiagen) for total DNA extraction, following the manufacturer’s protocol, with the addition of a 10 min incubation at 65°C after adding the lysis buffer and prior to the bead-beating step. DNA quantification was performed using the Qubit dsDNA HS Assay kit and Qubit 4 fluorometer (ThermoFisher).

### Shotgun virome library preparation and sequencing

Sequencing of all viromes was performed by the DNA Technologies and Expression Analysis Core at the University of California, Davis Genome Center (Davis, CA, USA). Metagenomic libraries were constructed with the DNA Hyper Prep kit (Kapa Biosystems-Roche, Basel, Switzerland) and sequenced (paired-end, 150 bp) using the NovaSeq S4 (Illumina, San Diego, CA, USA) platform to a requested depth of 10 Gbp per virome.

### Amplicon library preparation and sequencing

We performed 16S rRNA gene amplicon sequencing on all total DNA extractions with a dual-indexing strategy (49). We used the Platinum Hot Start PCR Master Mix (ThermoFisher) with the 515F/806R universal primer set for amplification of the V4 region of the 16S rRNA gene. We followed Earth Microbiome Project’s PCR protocol (50), which included an initial denaturation step at 94°C for 3 min, 35 cycles of 94°C for 45 s, 50°C for 60 s, and 72°C for 90 s, and a final extension step at 72°C for 10 min. Three of the samples had an undetectable total DNA yield, so only one attempt at amplification was tried for those, all of them failing to amplify (BSM25, USM26, BSM31). Two other samples had detectable total DNA yields but failed four attempts at amplification (BSM11, UDM33). We cleaned libraries with AmpureXP magnetic beads (Beckman-Coulter), quantified amplicons with a Qubit dsDNA HS Assay kit and Qubit 4 fluorometer (ThermoFisher), and then pooled all bead-purified products together in equimolar concentrations. The libraries were submitted to the DNA Technologies and Expression Analysis Core at the University of California, Davis Genome Center and sequenced (paired-end, 250 bp) using the MiSeq (Illumina) platform.

### Comparison to experimental heating

For our bioinformatics processing and data analysis, we also included soil viromes from a laboratory heating experiment performed on samples collected one year later (April 2022) at the same location of one of the control plots (plot U1), with samples subjected to different heat treatments in the laboratory (32). We included these samples to increase the number of recoverable vOTUs via read mapping (parameters described below) and for comparative analyses (parameters the same as for within-study comparisons, unless otherwise indicated).

### Virome bioinformatic processing

Viromes were processed as in our prior study (32). Briefly, raw reads were trimmed using Trimmomatic v0.39 (51) with a minimum q-score of 30 and a minimum read length of 50 bases. We used BBMap v39.01 (52) to remove PhiX sequences (k-mer size = 31 and hdist = 1). Each virome was assembled separately with MEGAHIT v1.2.9 (53) in the metalarge mode with a minimum contig length of 10,000 bp.

Assembled contigs were analyzed with VIBRANT v1.2.0 (54) with the -virome flag to identify viral contigs. We used CD-HIT v4.8.1 (55) to cluster all output viral contigs at 95% shared nucleotide identity and 85% alignment coverage (breadth). Quality-filtered raw reads from all viromes were then mapped against the representative set of vOTUs using Bowtie 2 v2.4.1 (56) in sensitive mode. Lastly, we used CoverM v0.5.0 (57) to quantify vOTU relative abundances in each sample from the mapped read coverage and generated a trimmed mean coverage table with ≥75% horizontal (breadth) coverage threshold for vOTU detection and a count table with absolute number of reads aligned to each contig for downstream analyses. Host taxonomy predictions for the entire set of dereplicated vOTUs was determined using iPHoP v1.1.0 (58), with the iPHoP_db_Sept21_rw database and default parameters (minimum cutoff score of 90) (**Supplementary Table 6**).

### 16S rRNA gene amplicon sequence bioinformatic processing

We used DADA2 v1.12.1 (59) to process 16S rRNA gene amplicon reads. We quality filtered reads with the filterAndTrim() function with the following parameters: truncLen = c(0,0), maxN = 0, maxEE = c(2,2), truncQ = 2, rm.phix = TRUE. Default settings were used to perform error rate inference, dereplication, denoising, read merging, and chimera removal. Amplicon sequence variants (ASVs) were assigned taxonomy via the DADA2 RDP classifier106, using the SILVA database v138.1 as reference (60).

### Statistical and ecological analyses

All ecological and statistical analyses were performed with R v4.3.0 (61). Input for all analyses of viral communities or of vOTUs (e.g., vOTU detection patterns across samples, vOTU relative abundances) was the trimmed mean coverage vOTU table. We normalized the vOTU table by using the decostand function (method = “total”, MARGIN = 2) from the Vegan package v2.6-6.1 (62) and removed singletons (vOTUs detected in only one sample). With a DADA2-generated relative abundance table for 16S rRNA gene amplicon sequence variants (ASVs), we filtered out mitochondria and chloroplasts and removed singletons (ASVs that only appeared in a single sample). To adjust for differences in the number of reads in each sample, we generated rarefaction curves using the Vegan function rarecurve (step = 100) and determined a rarefaction threshold, or read depth, for subsampling. The threshold was chosen as the number of reads that captured the highest number of different species for the vast majority of samples. We subsampled the reads to the chosen depth with the Vegan function rarefy. One sample (BSM27) was far below the threshold, so it was eliminated from further analysis. The following Vegan functions were used: vegdist (method = “bray”) to calculate Bray-Curtis dissimilarities, adonis2 to perform PERMANOVA, and mantel to perform Mantel tests.

We performed Kruskal-Wallis rank sum tests on means, followed by pairwise Wilcox tests if the Kruskal-Wallis test was significant, with the Bonferroni p-value adjustment method for multiple comparisons. We determined correlation of paired samples between two variables with the cor.test function (Spearman method). We used the cmdscale function to perform multidimensional scaling and retrieve true eigenvalues and calculate PCoA point values. We performed PCA analysis with the princomp function (base R).

We created upset plots with ComplexUpset v1.3.5, using vOTU or ASV relative abundance tables that were transformed into presence-absence tables (63). To determine the spatial distance between plots, we used the package geosphere v1.5-18 (64). We used the rcorr function from the package Hmisc v5.1-3 to determine correlations between relative abundances and chemical properties (65) and the pheatmap function from the package pheatmap v1.0.12 to visualize the correlations (66). Statistical analyses are documented in **Supplementary Tables 7-10**.

## RESULTS AND DISCUSSION

### Study design and dataset features

To determine the responses of soil dsDNA viral and prokaryotic communities to a prescribed burn, we designed a before-after control-impact (BACI) (67,68) field study with four treatment plots (burned) and two control plots (not burned) (**Figure 1A**). We sought to compare communities with and without burn treatment, both over time (before-after) and within timepoints (control-impact). In each of the six plots, we collected two replicate soil profiles with samples at two depths at 5 timepoints (14 and 3 days before and 7, 14, and 30 days after the prescribed burn, henceforth T1-5), for a total of 120 samples (**Figure 1B**).

We extracted DNA from DNase-treated viromes (91 viromes sequenced from the first four timepoints) and performed 16S rRNA gene amplicon sequencing on total DNA (115 samples amplified and sequenced from all timepoints). Fifteen viromes, or 12.5% of the total dataset had viromic DNA yields below detection limits (< 0.05 ng/µl, or < 0.49 ng/g soil), which was insufficient for sequencing (**Figure 2A**). Ten of those viromes were from T5 (30 days post-fire in soils approaching dryness, a condition previously shown to result in low-to-undetectable viromic yields (48)), with 8 from treatment plots and 2 from control plots. Due to low statistical power for treatment-control comparisons at T5, we decided against sequencing any of the viromes from that timepoint, resulting in a total of 29 viromes not sequenced from the full set of 120. We accounted for this ‘censored data’ (69) in our analysis of DNA yields (where the actual value for viromes with yields below detection limits was an unknown between 0 and 0.49 ng/g soil), and we report results from the sequenced viromes, with attempts to account for the missing data in our analyses and interpretations where possible.

**Figure 2.**
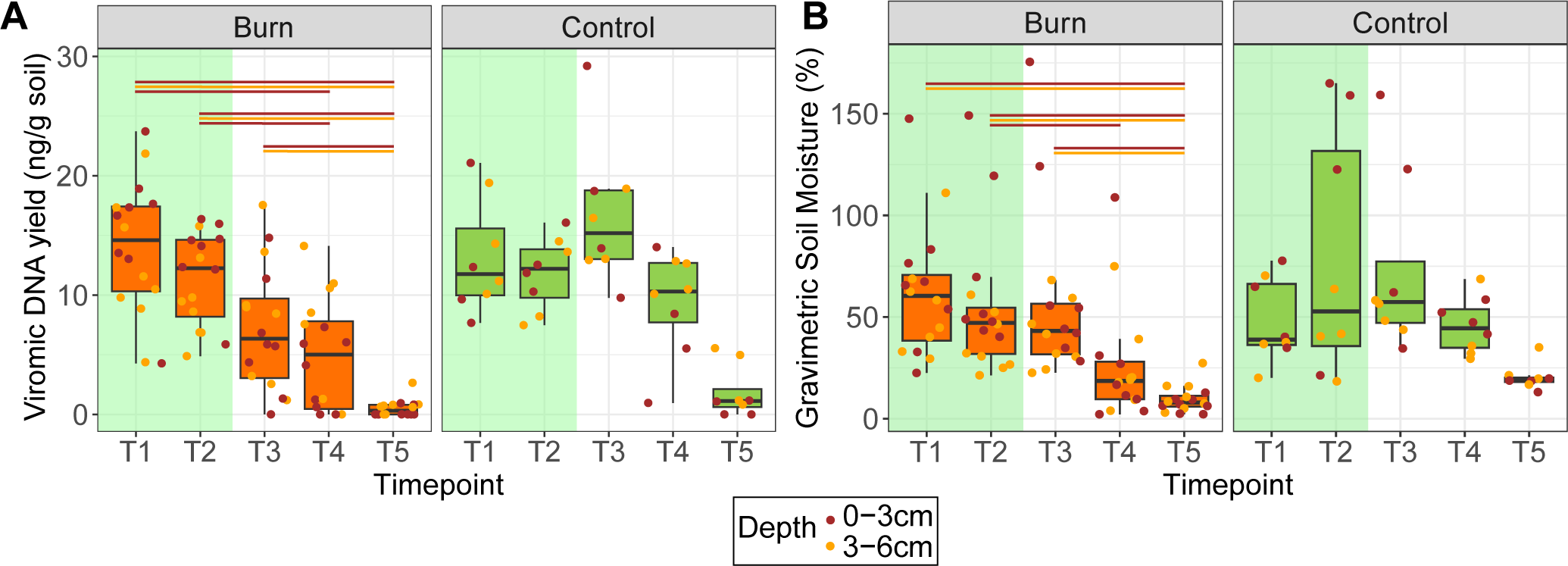
Viromic DNA yields and soil moisture content, **A.** DNA yields per gram of soil for each extracted virome at each timepoint, faceted by burn (left) and control (right) plots, with substitution of 0 ng/g soil for all viromes with yields that were below detection limits. Point colors indicate depth of sample. Box boundaries correspond to 25^th^ and 75^th^ percentiles, and whiskers extend to ±1.5x the interquartile range. Horizontal lines indicate whether there was a significant pairwise Wilcox test between timepoints and at which depth (color of lines). Panel background colors highlight pre-burn (green) and post-burn (white) timepoints. **B.** Gravimetric soil moisture percent for each sample at each timepoint, faceted by burn (left) and control (right). Colors, box and whisker parameters, and statistics are the same as in **A**. Timepoints T1-T5 correspond to 14 and 3 days before and 7, 14, and 30 days after the fire, respectively.

We assembled 327,006 contigs > 10 kbp that were identified as likely viruses using VIBRANT (54) and clustered them at 95% average nucleotide identity (ANI) with the 141,638 contigs assembled from our laboratory heating experiment on soils from the same site ((32), see methods) to a final count of 272,017 ‘species-level’ (70) vOTUs.

The total number of vOTUs in the current study was reduced to 198,402 after removing vOTUs only detected in the previous heating experiment and removing singletons (vOTUs only detected in one virome). From the 16S rRNA gene amplicon sequencing data, we recovered 9,210 ASVs from both studies, reduced to 4,136 ASVs after rarefying to a common depth of 18,500 reads per sample, removing singletons (ASVs that were detected in only one sample), and removing ASVs only detected in the heating experiment.

### Viromic DNA yields decreased significantly in post-burn samples

The first evidence that burn treatment had an impact on viral communities came from significant differences in viromic DNA yields, a proxy for viral particle abundances (32,48) (**Figure 2A**). Viromic DNA yields for both burn and control plots significantly changed over time at each depth (Kruskal-Wallis, p < 0.05), but a pairwise Wilcoxon test showed significant yield decreases in post-fire timepoints compared to pre-fire timepoints in the burn plots only (p.adj < 0.05) and no significant differences between any of the timepoints in the control plots. Surface soils (0-3 cm) had more significantly different pairwise comparisons between timepoints than soils from 3-6 cm (5 versus 3 of a total of 8 comparisons for each depth), potentially indicating greater impact from the fire in the surface soils, as would be expected according to previously observed surface-limited impacts of fire (25–28). At T3, the first timepoint after the fire (seven days post-burn), viromic DNA yields were significantly lower in burn versus control plots at 0-3 cm only, and at T5, they were significantly lower at 3-6 cm only. Otherwise, within each timepoint, there were no significant differences in viromic DNA yields between burn and control plots (which was expected for pre-fire timepoints, but not necessarily for post-fire timepoints) or between depths within each treatment. Since all viromes with DNA yields below detection limits are considered censored data with an unknown yield value between 0 and the lower detection threshold of 0.05 ng/µl (0.49 ng/g soil), we set those values to 0 ng/g soil (the lower bound) and, in a repeat analysis, to 0.49 ng/g soil (the upper bound), and there was no difference in the statistical results. Overall, results suggest that soil viral particle abundances decreased after the burn, particularly in near-surface soils within seven days of the burn, but that this response was ephemeral, with recovery of viral biomass (statistically indistinguishable from controls) within 14 days of the burn.

Soil moisture levels differed significantly over time at both depths for burn plots and at the surface depth for control plots (Kruskal-Wallis, p < 0.05). While most of our sampling occurred after recent precipitation events, T5 (30 days post-fire) did not (**Supplementary Figure 1A**). However, only in burn plots did pairwise Wilcoxon tests show a significant decrease in soil moisture at T5 compared to other time points (significant compared to T1-T3 at both depths, p.adj < 0.05) (**Figure 2B**), indicating that fire likely exacerbated drying conditions. Low-to-undetectable viromic DNA yields at T5 for samples from both control and burned plots are consistent with prior work suggesting lower viral biomass in dryer soils (48). Given that all but two viromes with DNA yields below detection limits were from post-fire treatment plots, along with significant decreases in both viromic DNA yields and soil moisture in burn plots only, we infer that the reduction of soil viral particle abundances can be attributed to burn treatment, both directly and in tandem with drying soil conditions.

### Spatial location better explained viral community composition than did burn treatment

When considering the full viromic dataset across plots, disturbance status (burned versus unburned), timepoints, and depths, plot (sample location) was the most significant factor driving viral community composition (explaining 17.32% (R^2^) of the total variation in the data, p = 0.001 by PERMANOVA) (**Figures 3A-B**). The remaining variables were significant (p < 0.05) but explained much lower percentages of the variation. Timepoint and disturbance status explained 4.95% and 1.73% of the total variation, respectively (both p = 0.001 by PERMANOVA), and depth explained 1.31% of the variation (p = 0.011 by PERMANOVA). A linear regression of pairwise spatial distances and viral community Bray-Curtis similarities across pre-fire samples (all plots, two timepoints) revealed a very weak but highly significant negative correlation at both T1 and T2 (p < 0.001 for both) (**Supplementary Figures 1B-C**), suggesting a distance-decay relationship, with communities farther apart more different than those closer together before any treatment had been applied. This relationship held after burn treatment for both treatment and control samples together (p < 0.001 for both T3 and T4) (**Supplementary Figures 1D-E**). Of the 198,402 vOTUs detected in more than one virome, the majority (57%) were detected in only one plot (**Figure 3C**), with only 1,100 vOTUs (0.55%) detected in all plots. The observed viral community compositional differences by plot, a subtle but significant distance-decay relationship across the sampled area, and the restricted detection of most vOTUs in single plots suggest that strong dispersal limitation and spatial structuring overlayed potential burn treatment effects at this site, consistent with other studies that have demonstrated strong spatial structuring of soil viral communities (40,71).

**Figure 3.**
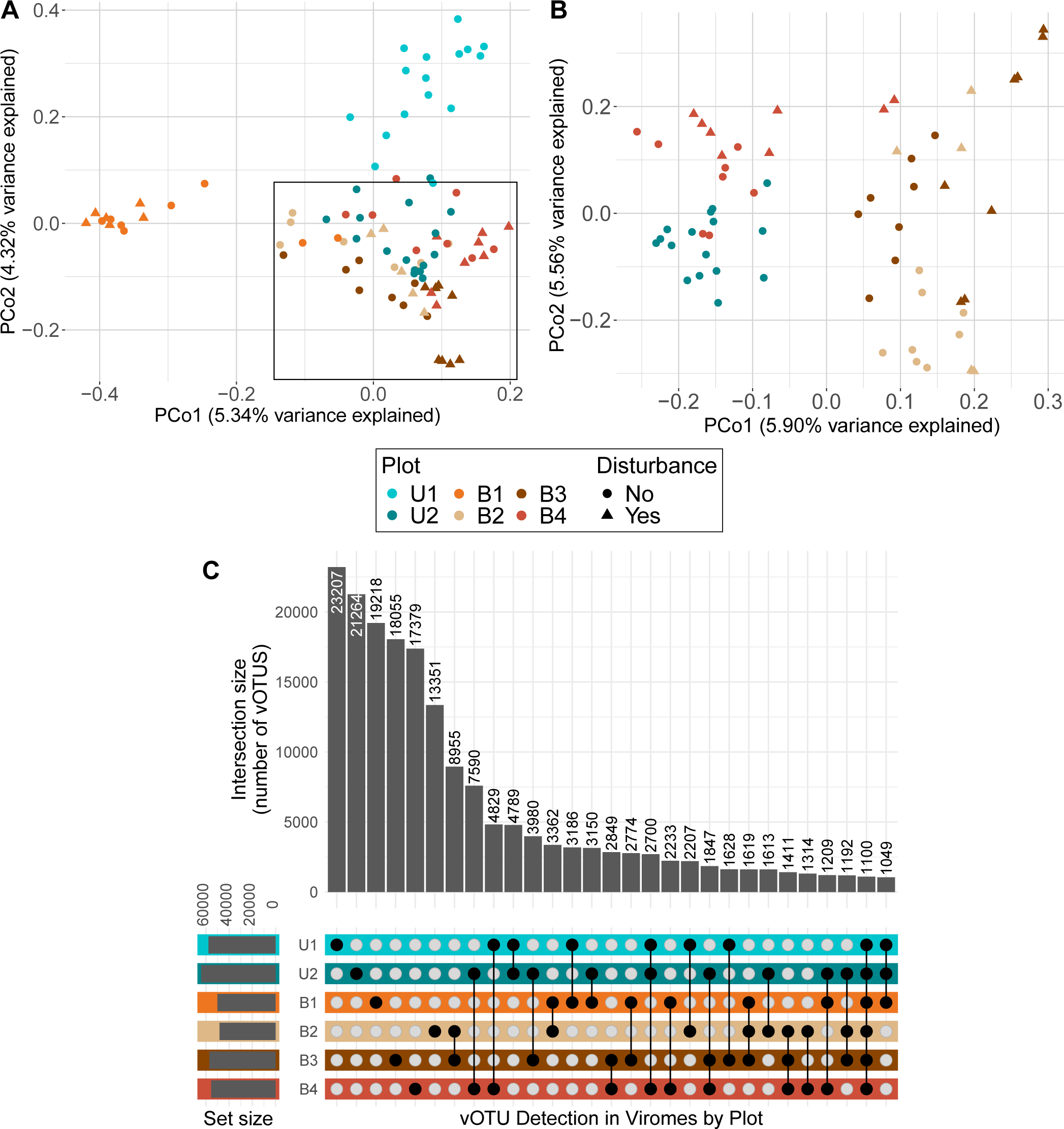
Spatial structuring of soil viral communities, **A/B.** Unconstrained analyses of principal coordinates (PCoA) performed on vOTU Bray-Curtis dissimilarities calculated across viromes, with colors indicating plot (location) and shapes indicating whether or not the sample experienced a burn disturbance. **B**. PCoA excluding samples from the two plots (U1 and B1) with viral communities that separated the most from the rest of the viromes (to better show trends for remaining viromes). Inset square in panel **A** approximately outlines panel **B** data. **C.** UpSet plot of vOTU detection patterns across sampled plots, with blackened circles indicating detection in a particular plot and intersection size indicating the number of unique vOTUs with a given detection pattern (found in one or more specific plots).

### Soil viral and prokaryotic community responses to fire were most evident at high burn severity

A comparison of soil chemical responses to burn treatment revealed a patchy, gradient-like response, ultimately resulting in the development of a chemistry-based burn severity index. We compared responses of individual environmental variables and, separately, the suite of all measured variables together between burned and unburned samples. For most measured properties, there was substantial variation across post-fire treatment (burned) samples when compared to both types of control samples (from pre-fire burn plots and unburned control plots) (**Supplementary Figures 2A-I**). The heterogeneity in measured chemical parameters was consistent with the heterogeneity in measured maximum temperatures achieved in each burned plot (a range from 29.7 to 474.8 °C at 2 cm depth and 17.5 to 363.6 at 6 cm depth), including differences on both sides of plot B1 (474.8°C on one side and 62.6°C on the other side, approximately 5 m apart) (**Supplementary Figure 3A**). In a principal components analysis (PCA) of ten burn-relevant chemical properties (soil pH, organic matter, soil moisture, organic phosphorus, inorganic phosphorus, inorganic nitrogen, organic nitrogen : inorganic nitrogen, magnesium, calcium, and potassium (22–24)), all unburned samples (control samples and burn plots pre-fire) clustered closely together, while the post-fire treatment samples had a large spread, with some clustering with the unburned samples and the remaining distributed along the PCA1 axis, which explained 47% percent of the variance in the data (**Figure 4A**). This variability is indicative of a heterogenous, or patchy, soil chemical response to fire, which is a known property of spring prescribed burns (39), and it suggests that accounting for different degrees of burn severity might reveal viral and prokaryotic community responses to fire that were masked in our overall comparisons of burned versus unburned samples above.

**Figure 4.**
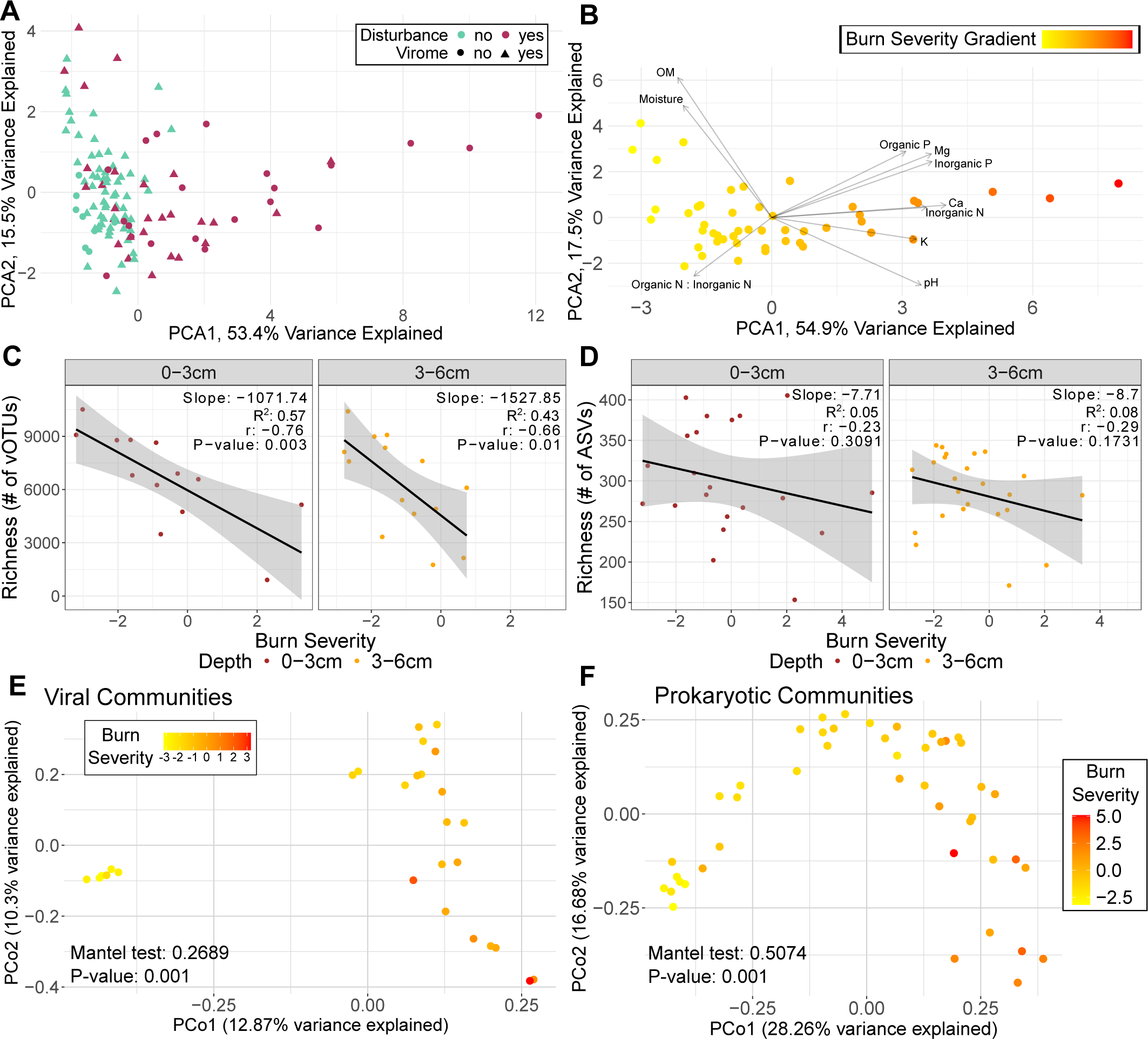
Burn severity as a function of soil chemistry and its effects on viral and prokaryotic communities, **A.** Principal component analysis (PCA) of z-transformed burn-relevant chemical properties (same properties as vector labels in **B**) for each sample. Colors indicate disturbance (burned or not burned) and shapes indicate whether a virome was sequenced for that sample. **B**. Subset of **A**, with only burned samples (burgundy points in **A**). Colors indicate the PCA1 scores (burn severity gradient), and vectors indicate the direction and magnitude of loadings (which are labeled at the end of their respective arrow, OM = organic matter). **C,D**. Richness of each burned virome (**C**) and 16S rRNA gene ASV profile (**D**) across the burn severity gradient. Trend lines display the least squares linear regression model. **E/F.** Principal Coordinates Analyses (PCoA) performed on viral community Bray-Curtis dissimilarities calculated only for viromes from burned plots (**E**) and prokaryotic community Bray-Curtis dissimilarities calculated only for 16S rRNA gene amplified samples from burned plots (**F**). Color gradient corresponds to chemistry PC1 scores (the burn severity index) from panel **B**.

Given the patchy soil chemical response to burn, we sought to create a burn severity index for our dataset that could be leveraged as an explanatory factor for differences in the viral and/or prokaryotic community responses across burned samples. We constructed a new PCA plot with the same subset of ten burn-relevant chemical properties described above, using burned samples alone (**Figure 4B**). The direction and magnitude of the loadings that contributed to the variation captured by PCA1 (shown as vectors in the PCA) aligned with known post-fire chemical changes (**Supplementary Figure 2J**), including an increase in pH and available nutrients and a decrease in organic matter and soil moisture (22–24). Therefore, we used the PCA1 scores from the PCA with burned samples only (**Figure 4B**) as a proxy for burn severity, assigning samples with lower values as less severely burned and samples with higher values as more severely burned. This allowed us to further interrogate impacts of burn treatment on viral and prokaryotic community composition beyond our binary burned vs. unburned comparisons and to assess differences in burn severity with depth.

We next investigated whether burn severity and/or burn treatment effects on viral and prokaryotic community composition were significantly different with depth in post-burn samples, as we hypothesized that there would be more substantial impacts at 0-3 cm compared to 3-6 cm, since heat typically penetrates only to about 3 cm, even in high severity fires (27,28). When comparing paired samples (samples from the same location but different depths), burn severity was higher at 0-3 cm than at 3-6 cm for 21 of the 24 pairs (87.5%) (**Supplementary Figure 3B**), which confirms that, at least in terms of chemical changes, surface soils generally had a greater response to fire than soils below 3 cm. However, pairwise Bray-Curtis dissimilarities between viromes from each depth within the same vertical soil profile were not significantly different (Kruskal-Wallis, p = 0.5835) for any interaction of time and treatment, meaning that viral community differences across sample sets were similar at both depths (**Supplementary Figure 3C**). Findings from a study one year post-fire, where viruses were mined from total metagenomes, did find significant differences of viral community composition by depth (33). In our study, we suspect that the lack of significant differences in viral community composition with depth in the burned samples is at least partially attributable to the missing virome sequencing data from most high-severity burned samples, due to insufficient viromic DNA yields (meaning, the impact of fire was so severe that it destroyed most of the viral particles). In the same analysis for prokaryotes, there was also a lack of significance for depth-resolved prokaryotic community composition among treatments (p = 0.3108).

Among burned samples, both viral and prokaryotic richness was negatively correlated with burn severity (**Figure 4C**). Viral richness had significant negative correlations at both depths, (p < 0.05), however, while prokaryotic richness had a negative correlation at both depths (which is consistent with other studies noting decreased prokaryotic richness after fire (19)), neither finding was significant (**Figure 4D**). Burn severity was significantly but weakly correlated with viral community structure (Mantel statistic r = 0.2689, p = 0.001) (**Figure 4E**) and more strongly correlated with prokaryotic community structure (Mantel statistic r = 0.5053, p = 0.001) (**Figure 4F**), consistent with previous findings of burn impacts on bacterial community composition after fire (72). Again, the missing virome data (largely from samples at the highest burn severities, **Figure 5B**) could explain why the correlation with burn severity was not as strong for viral as for prokaryotic communities, given that we had a more complete dataset at high burn severity for the prokaryotes. When considering viromic DNA yields, the likelihood of yields below detection limits (indicative of low viral particle abundances) significantly increased with increasing burn severity (binomial generalized linear model, p < 0.05), further reinforcing our interpretation that burn treatment was correlated with lower viral particle abundances. Decreased viromic DNA yields have previously been associated with soils with low moisture content (48) and soils that have experienced high temperatures (32), and here there was likely the interplay of both conditions that contributed to low viral particle abundances in more severely burned soils. Overall, results suggest a patchy microbiological and physicochemical response to prescribed fire that was better explained by a chemistry-based burn severity gradient than by simple comparisons of burned and unburned samples.

**Figure 5.**
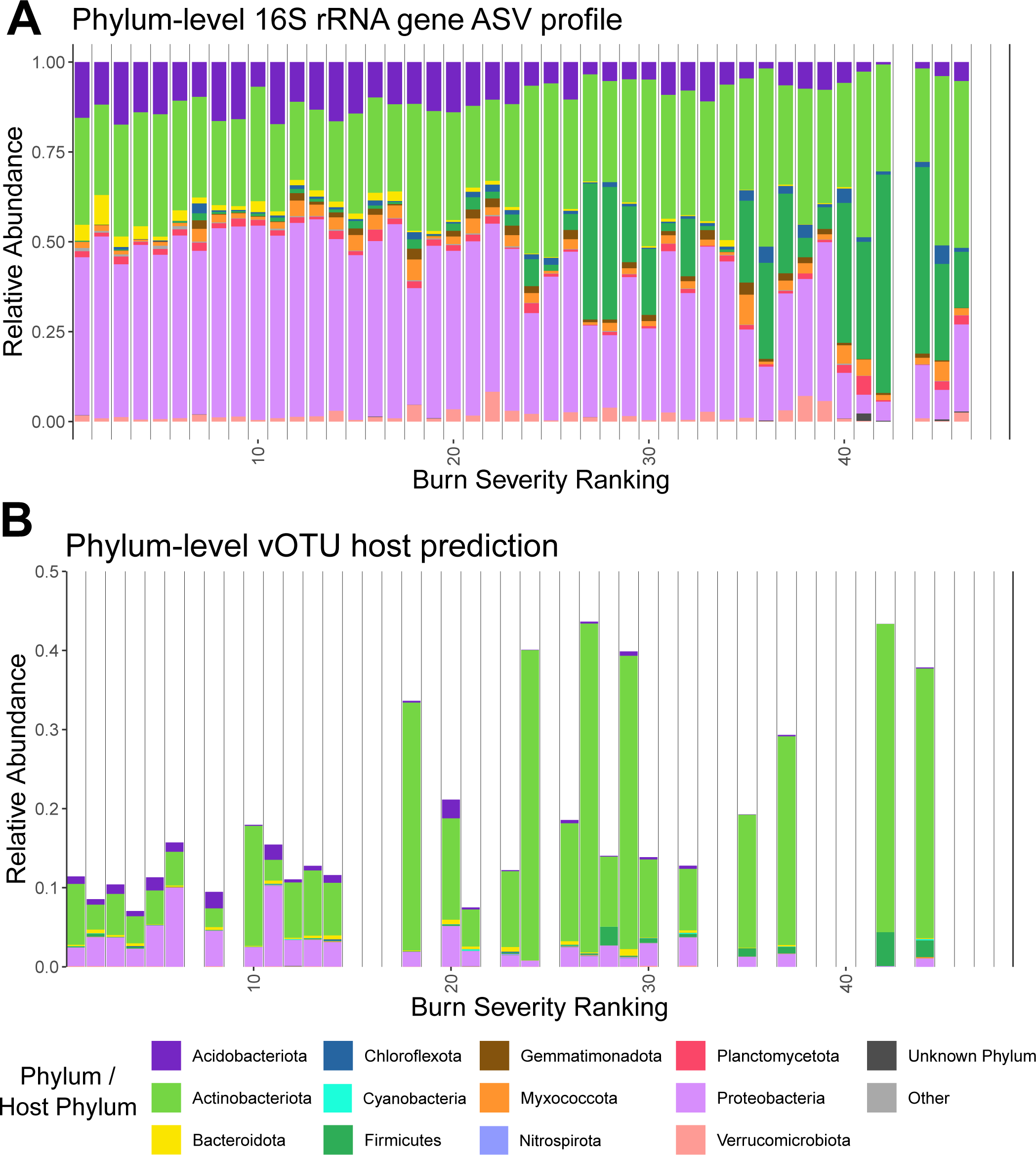
Dynamics along the burn severity gradient for prokaryotic taxa and for groups of viruses according to their predicted hosts, **A.** Phylum-level relative abundances in 16S rRNA gene profiles from burned samples. Each stacked bar is a sample. Missing bars indicate samples that did not amplify or meet the rarefaction threshold. Samples are in rank order of burn severity (low values on the left indicate low burn severity). The 10 most abundant phyla are colored, with all lower abundance phyla collapsed into “other.” **B.** Relative abundances of vOTUs with a host prediction in burned samples, grouped and colored by predicted host phylum (same legend as **A**). Host predictions were made with iPhoP ((58), see Methods). Each stacked bar is a virome, and missing bars indicate viromes that were not sequenced due to low or undetectable DNA yields. Viromes are in the same rank order of burn severity as for the prokaryotes in **A**.

### Viruses of the most fire-responsive prokaryotic taxa peaked in relative abundance at high burn severity

We next sought to identify burn-influenced shifts in the relative abundances of prokaryotic taxa and of the viruses predicted to infect them. When considering phylum-level prokaryotic taxonomy, trends were most clear when arranging samples along the burn severity gradient (**Figure 5A**), rather than by time or treatment group (**Supplementary Figures 4A-B**). The relative abundances of phyla Actinobacteriota, Chloroflexota, and Firmicutes had significant positive correlations with burn severity (Spearman’s, p < 0.05), with Firmicutes having a particularly high positive correlation (rho = 0.84). Stress-tolerant traits in these phyla, such as spore formation and performing well under drought conditions for members of the Actinobacteriota (73) and thermotolerance and spore formation in some members of both the Chloroflexota and Firmicutes (74,75), could explain their potential proliferation in burned soils. Additionally, Additionally, Actinobacteriota, Chloroflexota and aerobic members of Firmicutes (mostly Bacilli) are monoderm bacteria with thick cell walls which is thought to allow them to better resist drought stress (76). Increased relative abundances of these phyla, particularly Actinobacteria and Firmicutes, have also been seen in other post-fire studies (14,20,33) and in Blodgett forest soils (the same field site as this study) that had been heated to 60°C and 90°C (32). The relative abundances of phyla Acidobacteriota, Bacteroidota, and Proteobacteria had significant negative correlations with burn severity (Spearman’s rho > 0.75, p < 0.05). The reduction in Proteobacteria is counter to other post-fire studies, which have typically reported an increase in their relative abundance, with some Proteobacteria (e.g., in the genus *Massilia*) even being assigned as pyrophilous (13,14,77). A correlation analysis of phylum-level prokaryotic relative abundances and soil chemical properties across all samples (including non-burned controls) revealed that Firmicutes and Actinobacteriota were significantly correlated in the same direction with many of the same properties that explained variation captured in PC1 (our burn severity index) and that Acidobacteriota, Proteobacteria, and Bacteroidota were significantly negatively correlated (**Supplementary Figure 4C**).

Overall, phylum-level prokaryotic responses to fire were consistent with the emerging narrative from the rest of the study, reflecting a gradient-like response with burn severity, as opposed to a binary difference in taxa between burned and unburned samples.

Next, we investigated host predictions for our vOTUs to see whether viruses and their prokaryotic hosts exhibited similar dynamics in response to burn and/or across the burn severity range. Using iPhoP (58), this analysis was conducted at the host phylum level, since multiple predictions for the same vOTU had discrepancies at finer taxonomic scales. On a per-sample basis, we were able to predict a host for a minimum of 7% of vOTUs and a maximum of 44% of vOTUs, with an average of 20% and a significantly higher percentage of predictions in soils with higher burn severity (Spearman’s rho: 0.67, p < 0.05). As with prokaryotes, no clear trends in phylum-level host prediction were revealed based on timepoint or burn treatment (**Supplementary Figure 4E**), but with burn severity, clearer patterns emerged (**Figure 5B**). The relative abundances of viruses predicted to infect members of phylum Actinobacteriota or Firmicutes (two of the three prokaryotic phyla significantly positively correlated with burn severity) had a significant positive correlation with burn severity. Evidence of increased viral infections of Actinobacteriota was also reported in another post-fire study (33).

Conversely, the relative abundances of viruses with a predicted host in the phylum Acidobacteriota, Proteobacteriota, or Verrucomicrobiota (including two of the three prokaryotic phyla significantly negatively correlated with burn severity) had significant negative correlations with burn severity. A correlation analysis of relative abundances of viruses grouped by phylum-level host predictions and soil chemical properties across all samples (including non-burned controls) revealed that viruses predicted to infect Firmicutes, Actinobacteriota, and Myxococcota were significantly correlated in the same direction as most of the same properties that explained variation captured in PC1 (our burn severity index) (**Supplementary Figure 4D**). Together, these results indicate that viral population abundances responded to changes in host abundances and, either directly or indirectly, to soil chemical changes (e.g., via their own adapted traits, such as more burn-resistant structural elements, and/or by infecting adapted hosts) in a post-burn environment.

## CONCLUSIONS

This study interrogated soil viral community responses to a prescribed burn, revealing a heterogenous response aligned with burn severity. Viral communities were significantly spatially structured, with plot location explaining greater viral community compositional variation than burn treatment, depth, or time. Our empirically defined proxy for burn severity, generated from a suite of burn-relevant chemical properties measured precisely at each sampling location, was key to revealing trends in viral and prokaryotic community composition and taxon relative abundances that were otherwise obscured. When accounting for burn severity, viral and prokaryotic community richness and beta-diversity differed significantly according to the degree of burn. The relative abundances of four bacterial phyla significantly increased (Firmicutes and Actinobacteria, including known spore-formers) or decreased (Acidobacteriota and Proteobacteria) along the burn severity gradient, and the relative abundances of viruses predicted to infect these phyla responded in the same way as their putative hosts, suggesting increased viral predation of the most abundant (and, presumably, the most active) bacterial taxa. Increasing burn severity also corresponded with a greater number of low-to-undetectable viromic DNA yields, indicating reduced viral biomass with higher burn severity.

Our study reinforces the need to include robust measurements of treatment-related properties, such as soil chemistry, with sample-specific resolution for assessing prokaryotic and, particularly, viral community responses to inherently heterogenous disturbances like prescribed burning. Had we only considered coarse, binary ‘burned vs. unburned’ treatment conditions here, we would have lacked the resolution to explain the heterogenous soil viral and prokaryotic community responses to the prescribed burn, which were attributable to highly localized burn severity differences. Our analyses have provided evidence for a more nuanced soil community response to fire, not just to burn overall, but to the specific degree of burn severity experienced by each patch of soil, which was often different from the conditions of other nearby soils in the same fire.

## DATA AVAILABILITY

All raw sequences have been deposited in the NCBI Sequence Read Archive under the BioProject accession PRJNA1113186. The database of dereplicated vOTUs is available at https://zenodo.org/records/12562444. All scripts are available at https://github.com/seugeo/forestsoilviromes.

## ACKNOWLEDGEMENTS

We thank Patricia Lazicki for help with sample collection, staff at the Blodgett Forest Research Station (Ariel Thomson Roughton, Rob York, and Amy Mason) for advising on field sampling logistics, coordinating site visits, and coordinating the prescribed burn, the Traxler Lab at UC Berkeley (Monika Fischer, Neem Patel) for resource support, and Andrew Blandino (UC Davis Statistical Laboratory), Amisha Poret-Peterson (USDA Agricultural Research Service), Valerie Eviner (UC Davis Department of Plant Sciences), and Cole Smith for advice on statistical analysis. Thanks to Valerie Eviner and Jorge Rodrigues for helpful comments on a draft version of the manuscript. Funding for this work was provided by the U.S. Department of Energy (DOE), Office of Science, Office of Biological and Environmental Research (BER), Genomic Science Program, award number DE-SC0021198 (grant to JBE). SEG was also supported by the National Science Foundation Graduate Research Fellowship, the University of California Eugene Cota-Robles Fellowship, and the College of Agriculture and Environmental Sciences (UC Davis) Dean’s Circle Award. LSH was supported by the U.S. Department of Energy (DOE), Office of Science, Office of Biological and Environmental Research (BER), Genomic Science Program, award number DE-SC0023127 (grant to JBE), grant PI Sydney Glassman.

---

**Supplementary Figure 1.**
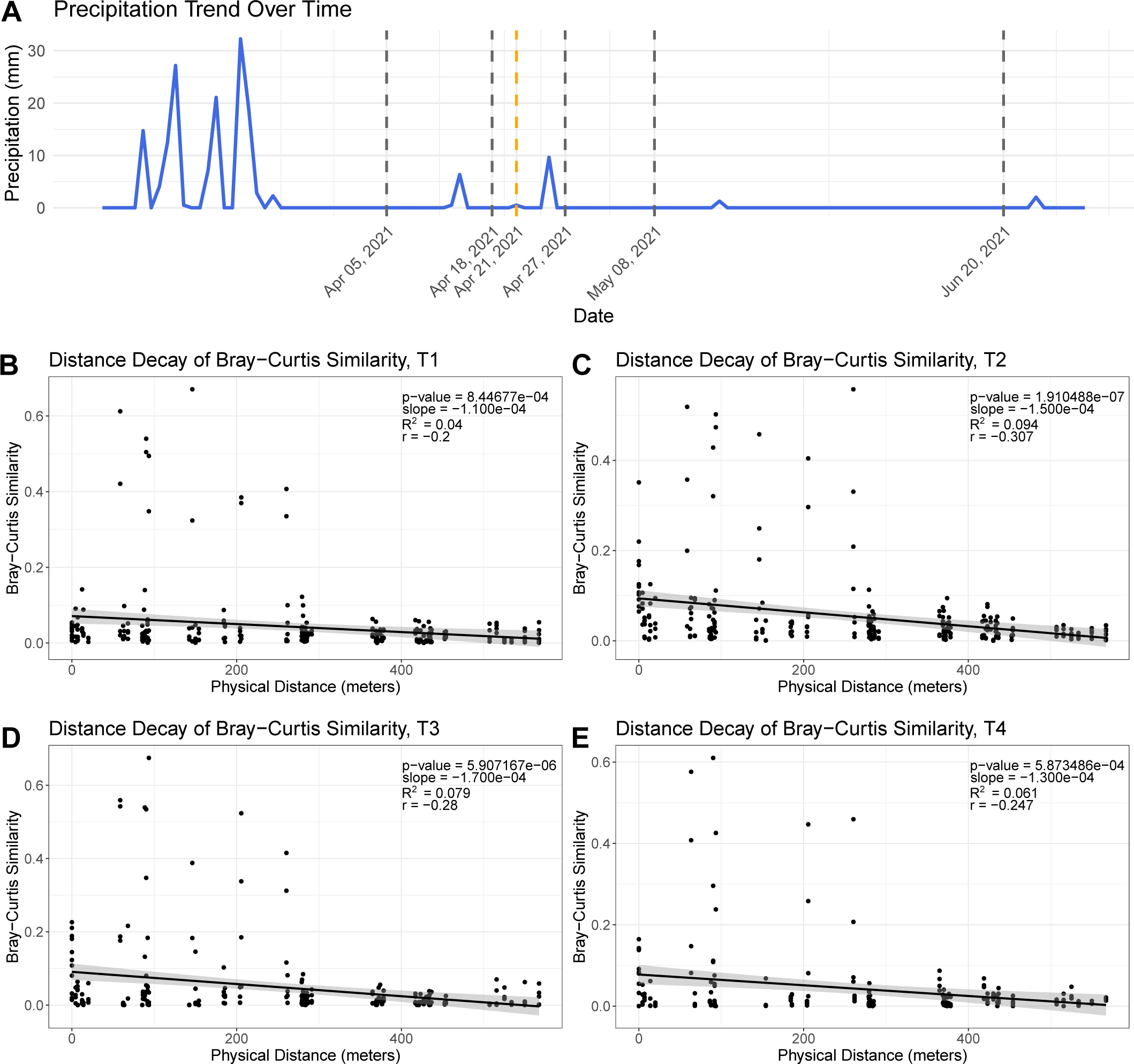
**A.** Amount of precipitation in millimeters measured each day between March 01, 2021 to June 30, 2021, with sampling dates highlighted with gray vertical lines and the prescribed burn date highlighted with an orange vertical line. **B-E.** Relationship between pairwise viral community Bray–Curtis similarity and pairwise spatial distance between samples, separated by timepoint to avoid time as a confounding factor [T1 (**B**), T2 (**C**), T3 (**D**), and T4 (**E**)]. Each point is a pair of samples. Trend lines display the least squares linear regression model. Inset statistics correspond to the Pearson’s correlation coefficient (r), R^2^, the linear regression slope, and the associated P value.

**Supplementary Figure 2.**
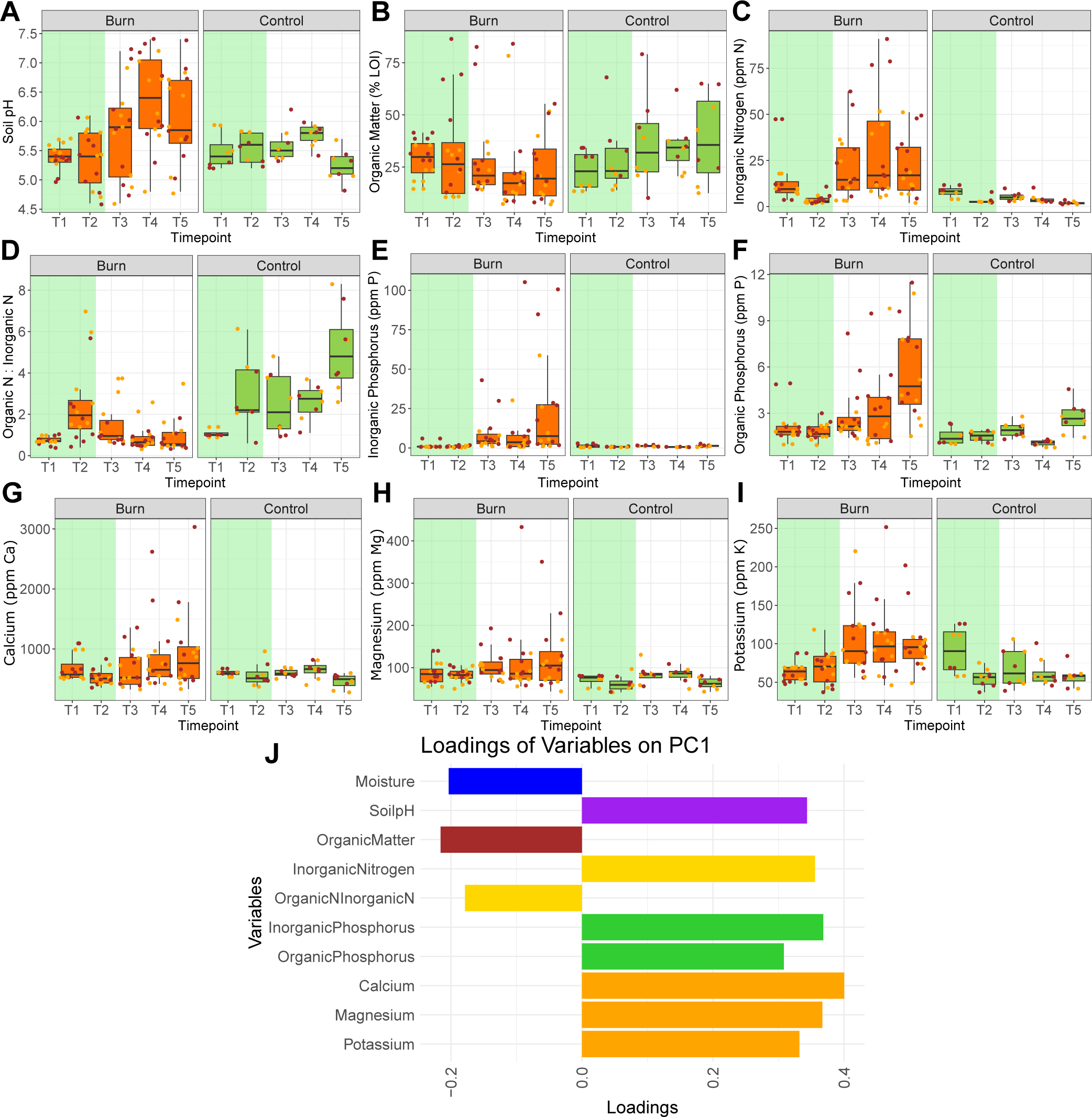
**A-I.** Burn-relevant soil chemical properties for each sample over time, faceted by burn (left) and control (right) plots. Point colors indicate depth of sample (brown 0-3 cm, orange 3-6 cm, same as in Figure 2). Box boundaries correspond to 25^th^ and 75^th^ percentiles, and whiskers extend to ±1.5x the interquartile range. For organic matter, LOI = loss on ignition (method). Panel background colors highlight pre-burn (green) and post-burn (white) timepoints. **J.** Principal component 1 (PC1) of the PCA in panel (**4B**) (the burn severity gradient) with bars representing the loadings along PC1 for each of the burn-relevant soil properties with different colors for either individual or groups of properties. Group colors: yellow = nitrogen properties, green = phosphorus properties, and orange = micronutrients.

**Supplementary Figure 3.**
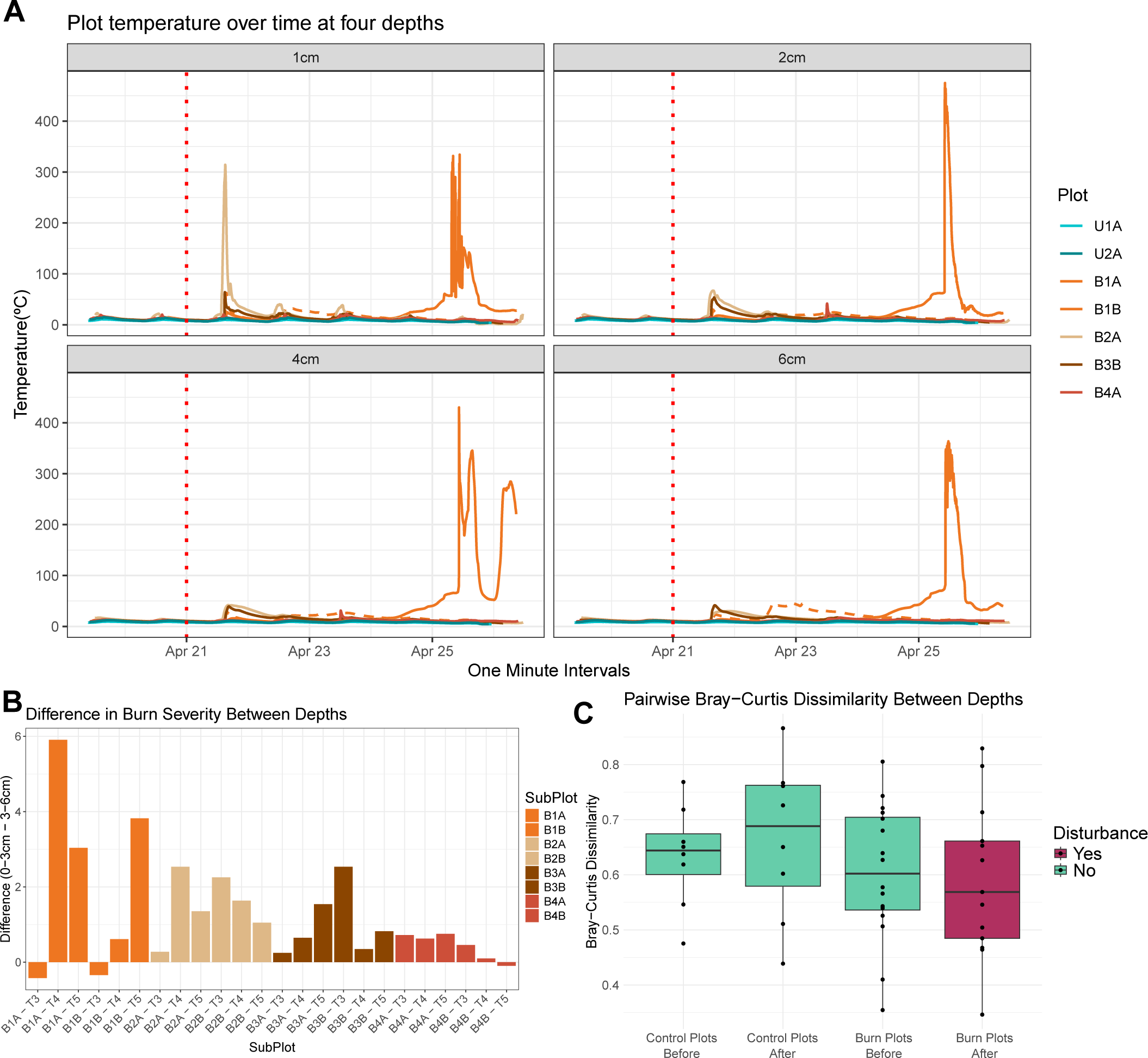
**A**. Time series temperature measurements from dataloggers placed at each plot (with two sets of data retrieved from plot B1) over the course of the prescribed burn. Figures are faceted by probe depth. Lines are colored by plot location. The red dotted vertical line approximated the beginning of the prescribed burn. **B**. Difference in burn severity between depths (upper depth severity minus lower depth severity) in burned plots, with each bar representing a subplot and timepoint and bars grouped and colored by plot. **C.** Boxplots representing the pairwise Bray-Curtis dissimilarity of viral communities between the two sample depths for each plot replicate at each timepoint. Box boundaries correspond to 25^th^ and 75^th^ percentiles, and whiskers extend to ±1.5x the interquartile range.

**Supplementary Figure 4.**
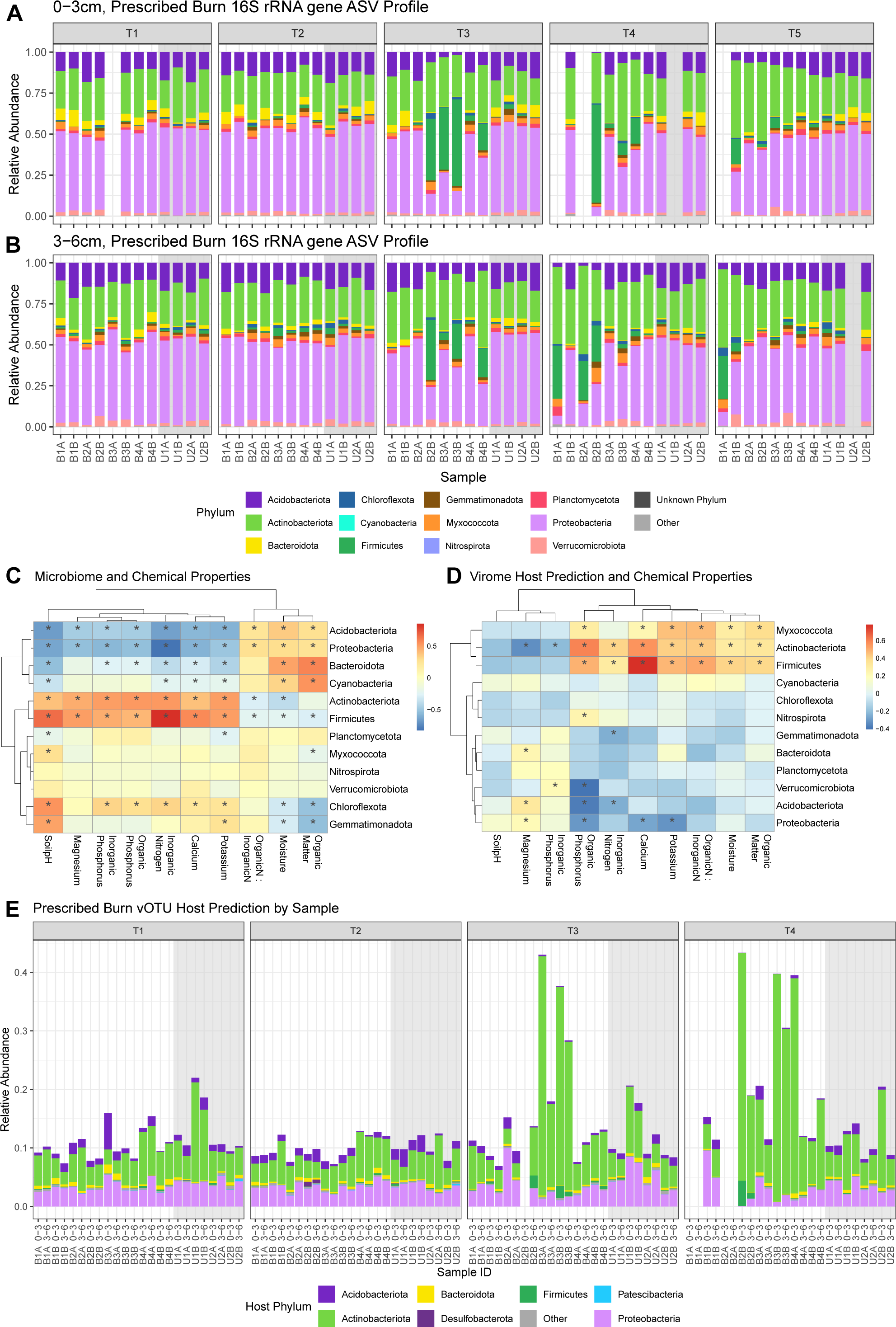
**A/B.** Phylum-level relative abundances in 16S rRNA gene profiles of all samples from 0-3 cm (**A**) and 3-6 cm (**B**). Each stacked bar is a sample, and missing bars indicate samples that did not amplify or meet the rarefaction threshold. Stacked bars are faceted by timepoint and grouped within each timepoint by plot. Panel background colors highlight control (gray) and burn (white) plots. The most abundant phyla are colored, with all other low abundance phyla collapsed into “other.” **C-D.** Hierarchical clustering and heatmap visualizing the correlation analysis of soil properties and relative abundances of prokaryotic phyla (**C**) or vOTU groups with a given phylum-level host prediction (**D**). Gradient of colors indicate Pearson’s correlation coefficient. Negative correlations correspond to dark blue and positive correlations correspond to dark red. Asterisks indicate a significant correlation (p < 0.05). **E.** Relative abundances of groups of vOTUs according to their phylum-level host predictions for all sequenced viromes. Each stacked bar is a virome. Missing bars indicate viromes that were not sequenced due to low or undetectable DNA yields. Stacked bars are faceted by timepoint and grouped within each timepoint by plot, with the sample depth indicated in the sample label. All other parameters are the same as (**A/B**).

